# Cell-type specific neuromodulation of excitatory and inhibitory neurons via muscarinic acetylcholine receptors in layer 4 of rat barrel cortex

**DOI:** 10.1101/2021.12.24.474122

**Authors:** Guanxiao Qi, Dirk Feldmeyer

**Affiliations:** Institute of Neuroscience and Medicine, INM-10, Reseach Centre Jülich, D-52425 Jülich, Germany; Department of Psychiatry, Psychotherapy and Psychsomatics, RWTH Aachen University, D-52074 Aachen, Germany; Jülich-Aachen Research Alliance-Brain, Translational Brain Medicine, D-52074 Aachen, Germany

**Keywords:** Acetylcholine, Muscarinic acetylcholine receptor, Nicotinic acetylcholine receptor, Layer 4, Barrel cortex

## Abstract

The neuromodulator acetylcholine (ACh) plays an important role in arousal, attention, vigilance, learning and memory. ACh is released during different behavioural states and affects the brain microcircuit by regulating neuronal and synaptic properties. Here, we investigated how a low concentration of ACh (30 µM) affects the intrinsic properties of electrophysiologically and morphologically identified excitatory and inhibitory neurons in layer 4 (L4) of rat barrel cortex. ACh altered the membrane potential of L4 neurons in a heterogeneous manner. Nearly all L4 regular spiking (RS) neurons responded to bath-application of ACh with a M4 muscarinic ACh receptor-mediated hyperpolarisation. In contrast, in the majority of L4 fast spiking (FS) and non-fast spiking (nFS) interneurons 30 µM ACh induced a depolarisation while the remainder showed a hyperpolarisation or no response. The ACh-induced depolarisation of L4 FS interneurons was much weaker than that in L4 nFS interneurons. There was no clear difference in the response to ACh for three morphological subtypes of L4 FS interneurons. However, in four morpho-electrophysiological subtypes of L4 nFS interneurons, VIP^+^-like interneurons showed the strongest ACh-induced depolarisation; occasionally, even action potential (AP) firing was elicited. The ACh-induced depolarisation in L4 FS interneurons was exclusively mediated by M1 muscarinic ACh receptors; in L4 nFS interneurons it was mainly mediated by M1 and/or M3/5 muscarinic ACh receptors. In a subset of L4 nFS interneurons, a co-operative activation of nicotinic ACh receptors was also observed. The present study demonstrates that low-concentrations of ACh affect the different L4 neurons types in a cell-type specific way. These effects result from a specific expression of different muscarinic and/or nicotinic ACh receptors on the somatodendritic compartments of L4 neurons. This suggests that even at low concentrations ACh may tune the excitability of L4 excitatory and inhibitory neurons and their synaptic microcircuits differentially depending on the behavioural state during which ACh is released.

## Introduction

Normal brain function relies on the participation of diverse neuromodulators such as the acetylcholine (ACh), noradrenaline, dopamine and serotonin. These neuromodulators are mainly released from different subcortical brain regions during different cognitive and behavioural states and affect neuronal microcircuits differently yet in a collaborative way. ACh plays a critical role in many cognitive functions including arousal, attention, vigilance, learning and memory *(Hasselmo 2006; Picciotto et al. 2012; Colangelo et al. 2019)*. While ACh is mainly released from axonal boutons of neurons located in the nucleus basalis of Meynert in the basal forebrain *(Mesulam et al. 1983; Zaborszky et al. 2015)*, it may also be co-released from neocortical choline acetyltransferase (ChAT)-expressing/vasoactive intestinal peptide (VIP)-positive interneurons together with the inhibitory transmitter GABA and/or VIP *(Obermayer et al. 2019; Granger et al. 2020)*. ACh effects are mediated by two different types of receptors, the G-protein-coupled muscarinic ACh receptors (mAChRs) and the ionotropic nicotinic ACh receptors (nAChRs). In the neocortex, both receptor types show layer-specific distributions and effects *(Obermayer et al. 2017; Radnikow and Feldmeyer 2018)*. In general, ACh increases the excitability of pyramidal cells located in different cortical layers by activating both nAChRs and mAChRs *(Gulledge et al. 2007; Zolles et al. 2009; Bailey et al. 2010; Tian et al. 2014; Hay et al. 2016; Yang et al. 2020; Patel et al. 2021)*. In a minor fraction of deep L2/3 and a subset of L5/6 pyramidal cells, ACh induces an initial small and transient hyperpolarisation followed by a sustained depolarisation mediated by muscarinic M1/3 mAChRs *(Gulledge and Stuart 2005; Gulledge et al. 2007; Eggermann and Feldmeyer 2009; Patel et al. 2021)*. In contrast, excitatory neurons located in layer 4 are persistently hyperpolarised by ACh activating M4 mAChRs *(Eggermann and Feldmeyer 2009; Dasgupta et al. 2018)*. A similar ACh effect was also found in L6A corticocortical neurons *(Yang et al. 2020)*.

Cholinergic effects on GABAergic inhibitory interneurons are heterogenous and dependent on interneuron subtypes *(Bacci et al. 2005; Muñoz and Rudy 2014)*. Cortical interneurons can be broadly divided into two large groups according to their firing patterns, i.e. fast spiking (FS) and non-FS (nFS) interneurons. ACh induces a depolarisation in the majority of nFS interneurons (e.g. somatostatin-expressing (SST+) adapting firing, VIP+ irregular spiking interneurons) via the activation of nAChRs and/or mAChRs but induces a hyperpolarisation in others such as cholecystokinin-expressing (CCK+) regular spiking interneurons *(Kawaguchi 1997; Gulledge et al. 2007)*. Whether FS interneurons show ACh effects is still a matter of debate *(Kawaguchi 1997; Xiang et al. 1998; Gulledge et al. 2007; Kruglikov and Rudy 2008; Chen et al. 2015)*.

The ACh response of a neuron depends on the concentration and the speed and spatial profile of application. In the majority of studies, a high concentration of ACh (∼1 mM) was applied locally through a puff pipette; this approach reveals predominantly the nicotinic ACh response but largely obscures any muscarinic ACh effects. Furthermore, a puff-application mimics (to some extent) phasic ACh release on a short time scale (within a few ms) but does not simulate tonic, non-synaptic ACh release into the extracellular space, the so-called ‘volume transmission’ *(Fuxe and Borroto-Escuela 2016)*.

In sensory cortices, layer 4 neurons receive direct thalamocortical input and distribute intracortical excitation and inhibition to other cortical layers. While the neuronal composition and synaptic connectivity of layer 4 have been studied extensively (Feldmeyer et al. 1999; Gibson et al. 1999; Lubke et al. 2000; Beierlein et al. 2003; Xu et al. 2013; Koelbl et al. 2015; Emmenegger et al. 2018; Scala et al. 2019) a comprehensive study on their modulation by ACh or other neuromodulators is still lacking. Here, we investigated how low concentrations of ACh affect the intrinsic properties of different L4 neuron types and subtypes in acute brain slices using patch-clamp recordings and bath-application of cholinergic agonists and antagonists. To reveal the cell-type specific effects of ACh, L4 neurons were classified into three electrophysiological types and ten electro-morphological subtypes as identified previously (Feldmeyer et al. 1999; Staiger et al. 2004; Koelbl et al. 2015; Emmenegger et al. 2018). We found that neuromodulation by mAChRs is a common property of all L4 neurons but is highly cell-specific. In addition, some L4 nFS interneuron types low concentrations of ACh evoked a strong superthreshold depolarisation mediated by coincident activation of both mAChRs and nAChRs suggesting that they are central in cholinergic modulation.

## Materials and Methods

All experimental procedures involving animals were performed in accordance with the guidelines of the Federation of European Laboratory Animal Science Association (FELASA), the EU Directive 2010/63/EU, and the German animal welfare law.

### Slice preparation

In this study, Wistar rats (Charles River, either sex) aged 18–33 postnatal days (P18-P33) were anaesthetized with isoflurane at a concentration < 0.1% and decapitated. The brain was quickly removed and placed in an ice-cold modified artificial cerebrospinal fluid (ACSF) containing a high Mg^2+^ and a low Ca^2+^ concentration (4 mM MgCl_2_ and 1 mM CaCl_2_), other components are same to that in the perfusion ACSF as described below, to reduce potentially excitotoxic synaptic transmission during slicing. In order to maintain adequate oxygenation and a physiological pH level, the solution was constantly bubbled with carbogen gas (95% O_2_ and 5% CO_2_). Thalamocortical slices *(Feldmeyer et al. 1999; Qi et al. 2017)* were cut at 350 µm thickness using a Leica VT1000S vibrating blade microtome and then transferred to an incubation chamber containing preparation solution for a recovery period of at least 30 minutes at room temperature before being transferred to the recording chamber. For relatively older animals (> P21), after cutting slices were transferred to a holding chamber placed in a water bath at 35 °C for 30 min and then, the water bath was allowed to gradually cool down to the room temperature.

### Solution

During recordings, slices were continuously superfused (perfusion speed ∼5 ml/min) with ACSF containing (in mM): 125 NaCl, 2.5 KCl, 1.25 NaH_2_PO_4_, 1 MgCl_2_, 2 CaCl_2_, 25 NaHCO_3_, 25 D-glucose, 3 mho-inositol, 2 sodium pyruvate and 0.4 ascorbic acid, bubbled with carbogen gas and maintained at 30-33 °C. Patch pipettes (5-8 MΩ) were pulled from thick-wall borosilicate glass capillaries and filled with an internal solution containing (in mM): 135 K-gluconate, 4 KCl, 10 HEPES, 10 phosphocreatine, 4 Mg-ATP, and 0.3 GTP (pH 7.4 with KOH, 290-300 mOsm). Biocytin at a concentration of 5 mg/ml was added to the internal solution in order to stain patched neurons after recordings.

### Electrophysiological recording and analysis

The slices and neurons were visualised using an upright microscope equipped with an infrared differential interference contrast (IR-DIC) optics. The barrels can be identified in layer 4 as dark stripes with light ‘hollows’ at low magnification (4x objective) and were visible in 6-8 consecutive slices. Neurons located inside the barrels were randomly selected for recordings. When being visualised at high magnification (40x magnification), putative excitatory neurons have ovoid-shape somata without obvious apical dendrites and putative interneurons have enlarged oval somata. They could also be differentiated by their intrinsic action potential (AP) firing patterns during recording and by their morphological appearances thereafter. Whole-cell patch clamp recordings were made using an EPC10 amplifier (HEKA, Lambrecht, Germany). Signals were sampled at 10 kHz, filtered at 2.9 kHz using Patchmaster software (HEKA), and later analyzed off-line using Igor Pro software (Wavemetrics, USA).

Custom-written macros in Igor Pro 6 (WaveMetrics, Lake Oswego, USA) were used to analyse the recorded electrophysiological signals. Passive and active AP firing properties were assessed by eliciting a series of 1 s current pulses under current clamp configuration. Neurons with a series resistance exceeding 40 MΩ or with a depolarized membrane potential (> -55 mV) after rupturing the cell membrane were excluded from analysis. The resting membrane potential (V_rest_) was recorded immediately after establishing the whole-cell recording configuration. Other passive membrane properties such as the input resistance R_in_, membrane time constant τ_m_, voltage sag were measured from membrane potential (V_m_) traces induced by a series of hyper- and depolarizing subthreshold current pulses. Single AP properties such as the AP threshold, amplitude, half-width, afterhyperpolarisation (AHP) amplitude were measured for the first spike elicited by a rheobase current step. Repetitive firing properties such as the maximum firing frequency, slope of frequency-current curve were measured. The description of most electrophysiological parameters for data analysis have been given previously *(Emmenegger et al. 2018)*.

### Drug application and analysis

Acetylcholine (ACh, 30 µM) was applied through the perfusion system. Atropine (ATRO, 200 nM), mecamylamine (MEC, 10 µM), tropicamide (TRO, 1 µM), pirenzepine (PIR, 0.5 µM), dihydro-ß - erythroidine (DHßE, 10 µM), TTX (0.5 µM) and the cocktail of synaptic blockers including CNQX (10 µM), D-AP5 (50 µM), gabazine (10 µM) were all bath-applied; drugs were purchased from Sigma-Aldrich or Tocris. During recordings, a 3 min stable baseline with a Vm fluctuation < 1 mV was recorded before applying the drug via the perfusion system. The change in Vm was calculated as the difference between the maximum Vm deflection (positive or negative) after drug application and the baseline. To avoid a misclassification of the Vm change because of background Vm fluctuation, we set a threshold of ± 0.5 mV so that a Vm change ≤ 0.5 mV during drug application is considered to be no response.

### Immunohistochemical staining

Slices were fixed after electrophysiological recordings with 4% paraformaldehyde (PFA) in 100 mM phosphate buffered saline (PBS) for at least 24 h at 4°C. To recover the morphology of biocytin-filled neurons, slices were rinsed several times in 100 mM PBS and then treated with 1% H_2_O_2_ in PBS for about 20 min in order to reduce any endogenous peroxidase activity. Slices were rinsed repeatedly with PBS and then incubated in 1% avidin-biotinylated horseradish peroxidase (Vector ABC staining kit, Vector Lab. Inc., Burlingame, USA) containing 0.1% Triton X-100 for 1 h at room temperature. The reaction was catalysed using 0.5 mg/ml 3,3-diaminobenzidine (DAB; Sigma-Aldrich, St.Louis, Mo, USA) as a chromogen. Slices were then rinsed with 100 mM PBS, followed by slow dehydration with ethanol in increasing concentrations and finally in xylene for 2– 4 h. After that, slices were embedded using Eukitt medium (Otto Kindler GmbH, Freiburg, Germany).

### Morphological reconstruction and analysis

Computer-assisted morphological 3D reconstructions of neurons were made using the NEUROLUCIDA® software (MicroBrightField, Williston, VT, USA) and Olympus BV61 microscopy at 1000x magnification (100x objective, 10x eyepiece). Neurons were selected for reconstruction based on the quality of biocytin labelling when background staining was minimal. The cell body, dendritic and axonal branches were reconstructed manually under constant visual inspection to detect thin and small collaterals. Cytoarchitectonic landmarks such as barrels in the primary somatosensory cortex and layer borders, pial surface and white matter were delineated during reconstructions at a low magnification (4x objective). The position of soma and layers were confirmed by superimposing the DIC images taken during the recording. Tissue shrinkage was corrected using correction factors of 1.1 in the x–y direction and 2.1 in the z direction *(Marx et al. 2012)*.

### Statistical analysis

For all data, the mean ± s.d. is given. Statistical comparisons among multiple groups were done using a Kruskal-Wallis test followed by a Dunn-Holland-Wolfe non-parametric multiple comparison test. Wilcoxon Mann-Whitney U test was performed to assess significant differences between individual groups. To assess the differences between two paired groups under different pharmacological conditions, Wilcoxon signed-rank test was performed. Correlation analysis was performed by calculating Pearson’s linear correlation coefficients. Statistical significance was set at p < 0.05, n indicates the number of neurons analysed. To prepare box plots for dataset with n > 10, the web application PlotsOfData was used (https://huygens.science.uva.nl/PlotsOfData/; Postma and Goedhart 2019).

## Results

We performed single-cell patch-clamp recordings in combination with biocytin fillings in acute brain slices to characterise the modulatory effect of ACh on the intrinsic properties of L4 neurons in the primary somatosensory (barrel) cortex of rats. In total, we have tested the effects of a low concentration of ACh (30 µM) on 108 L4 excitatory and inhibitory neurons. The ACh responses of L4 neurons was highly diverse depending on their electrophysiological and morphological identities.

### Electro-morphological classification of L4 neurons

Based on their electrophysiological characteristics, L4 neurons can be broadly classified as regular spiking (RS) excitatory neurons, FS and nFS inhibitory interneurons (**Fig. 1A**). Three L4 neuron types can be easily differentiated by only three electrophysiological parameters, i.e. the maximum firing frequency, AP half-width and the AHP amplitude (**Fig. 1B,C**). L4 RS neurons show a regular spiking firing pattern with a prominent spike frequency adaptation during a 1 s depolarising pulse (**Fig. 1B,C**). In contrast, L4 FS interneurons show a high-frequency firing pattern without obvious spike frequency adaptation. L4 interneurons of the nFS type show heterogeneous firing patterns including adaptive spiking, irregular spiking, late spiking etc.

**Figure 1.**
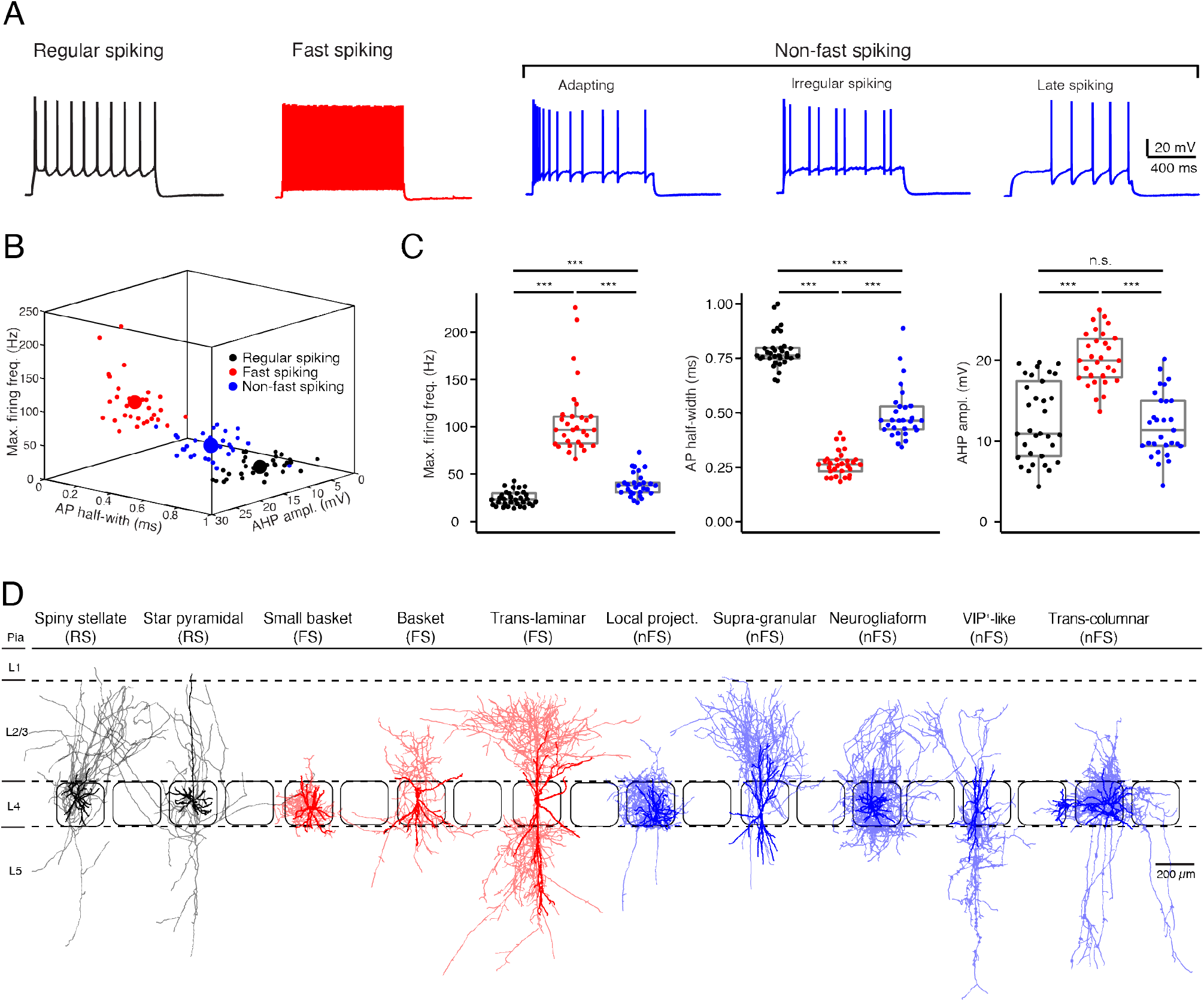
Electro-morphological classification of L4 neurons in the rat barrel cortex. (A) Representative firing patterns of L4 regular spiking (RS, black), fast spiking (FS, red) and non-fast spiking (nFS, blue) neurons. (B) Electrophysiological differentiation of L4 RS, FS and nFS neurons using the maximal firing frequency, AP half-width and the AHP amplitude. Mean and individual values are shown given by large and small dots, respectively. (C) Box plots of three electrophysiological parameters for L4 RS, FS and nFS neurons. P value was calculated using the non-parametric Wilcoxon-Mann-Whitney two-sample rank test. * p < 0.05, * * p < 0.01, * * * p < 0.001. (D) Morphological sub-classification of L4 RS, FS and nFS neurons. Soma and dendrites, opaque colour; axons, half-transparent colour.

In addition to their electrophysiological diversity, L4 neurons show highly distinct dendritic and in particular axonal morphologies (**Fig. 1D**). L4 excitatory neurons fall into two main groups, spiny stellate neurons (SSNs) without an obvious apical dendrite and star pyramidal cells (SPCs) *(Feldmeyer et al. 1999; Lubke et al. 2000)*; but see *(Staiger et al. 2004)*. Their axons originate from the soma or the initial part of one basal dendrite and project locally in layer 4 and to supra- and infragranular layers. Dendrites of L4 interneurons are aspiny or sparsely spiny and exhibit e.g. small to large multipolar, bipolar, or bitufted orientation patterns. Their axons project either locally in layer 4 and/or to supra- and/or infragranular layers in the vertical direction and/or to neighbouring columns in the horizontal direction. In previous studies, we have classified rat L4 FS interneurons as small basket cells (sBCs), basket cells (BCs) and translaminar cells (TLCs) *(Koelbl et al. 2015)* and L4 nFS interneurons as local-projecting (LP; non-Martinotti cell-like), supragranular-projecting (SP; Martinotti cell-like), neurogliaform (NGF), VIP^+^-like (VIP) and transcolumnar-projecting, interneurons *(Emmenegger et al. 2018)* (**Fig. 1D**).

### ACh at low concentrations induces diverse changes in the Vm of L4 neurons

We bath-applied 30 µM ACh while monitoring changes in the V_m_ of L4 neurons under current-clamp conditions. Of 44 L4 RS neurons, 42 showed a V_m_ hyperpolarisation; only two showed no change (**Fig. 2A,D**). On average, ACh-induced V_m_ change in L4 RS neurons was -2.8 ± 1.4 mV (n = 44) (**Fig. 3A**). Of 33 L4 FS interneurons, 25 neurons showed a weak but significant depolarisation of the V_m_, four a weak hyperpolarisation and another four no change (**Fig. 2B,E**). On average, the ACh application resulted in a change in V_m_ in L4 FS interneurons was 0.9 ± 1.5 mV (n = 33) (**Fig. 3A**). Of 31 L4 nFS interneurons, 29 neurons showed a strong depolarisation of the V_m_ and two a hyperpolarisation (**Fig. 2C,F**). For the majority of L4 nFS interneurons (29 out of 31), the ACh-induced depolarisation was subthreshold. In a small fraction of L4 nFS interneurons (4 out of 31), ACh application evoked a suprathreshold depolarisation so that spontaneous AP firing was initiated. On average, the ACh-induced change in V_m_ in L4 nFS interneurons was 5.2 ± 5.8 mV (n = 31) (**Fig. 3A**). Note that the ACh-induced V_m_ changes were fully reversible by bath application of control ACSF (**Fig. 2 A-C**). To examine whether these ACh-induced changes in V_m_ resulted from a direct effect on the neuronal excitability or were caused indirectly by altering the activity of local synaptic microcircuits, a cocktail of synaptic blockers comprising CNQX (10 µM), D-AP5 (50 µM) and gabazine (10 µM) was applied before ACh. There is no difference in the V_m_ change elicited by ACh in the absence and in the presence of synaptic blockers (**Fig. S1**). However, a clear decrease in background noise of V_m_ was observed in the presence of synaptic blockers (**Fig. S1**). A correlation analysis between the change in V_m_ and the age of the animal, V_rest_ and input resistance (R_in_) demonstrated that there is no clear age-dependence of the ACh effect on V_m_ for any of the three L4 neuron types (**Fig. S2A**); a significant negative correlation was found between the V_m_ change and V_rest_ for L4 RS neurons (r = -0.53, p = 1.8 × 10_-4_) and L4 nFS interneurons (r = -0.48, p = 5.9 × 10_-3_) (**Fig. S2B**). Furthermore, for L4 nFS interneurons, a significant positive correlation was found between the ACh-induced change in V_m_ and R_in_ (r = 0.77, p = 1.6 × 10_-7_; **Fig. S2C**), i.e. L4 nFS interneurons with higher R_in_ showed a larger depolarisation.

**Figure 2.**
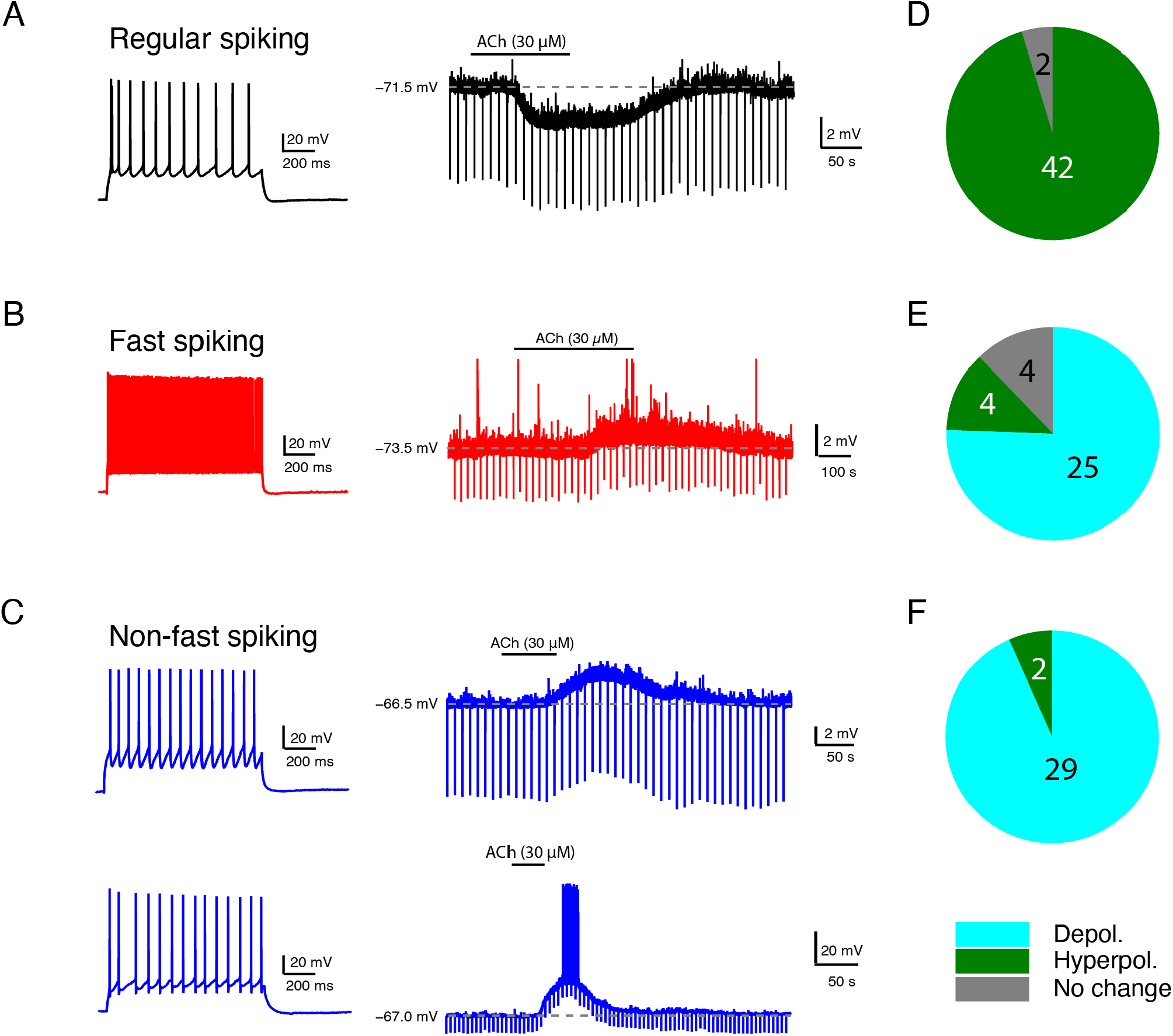
Low concentration of ACh induces diverse Vm changes in L4 RS, FS and nFS neurons. (A) An example recording of the time course of ACh-induced Vm change in a L4 RS neuron. (B) Same as (A) but for a L4 FS interneuron. (C) Same as (A,B) but for two different L4 nFS interneurons: top, sub-threshold depolarisation; bottom, supra-threshold depolarisation. (D) -(F) Pie charts summarising the ACh-induced changes in V_m_ in L4 RS (top), FS (middle) and nFS (bottom) neurons.

**Figure 3.**
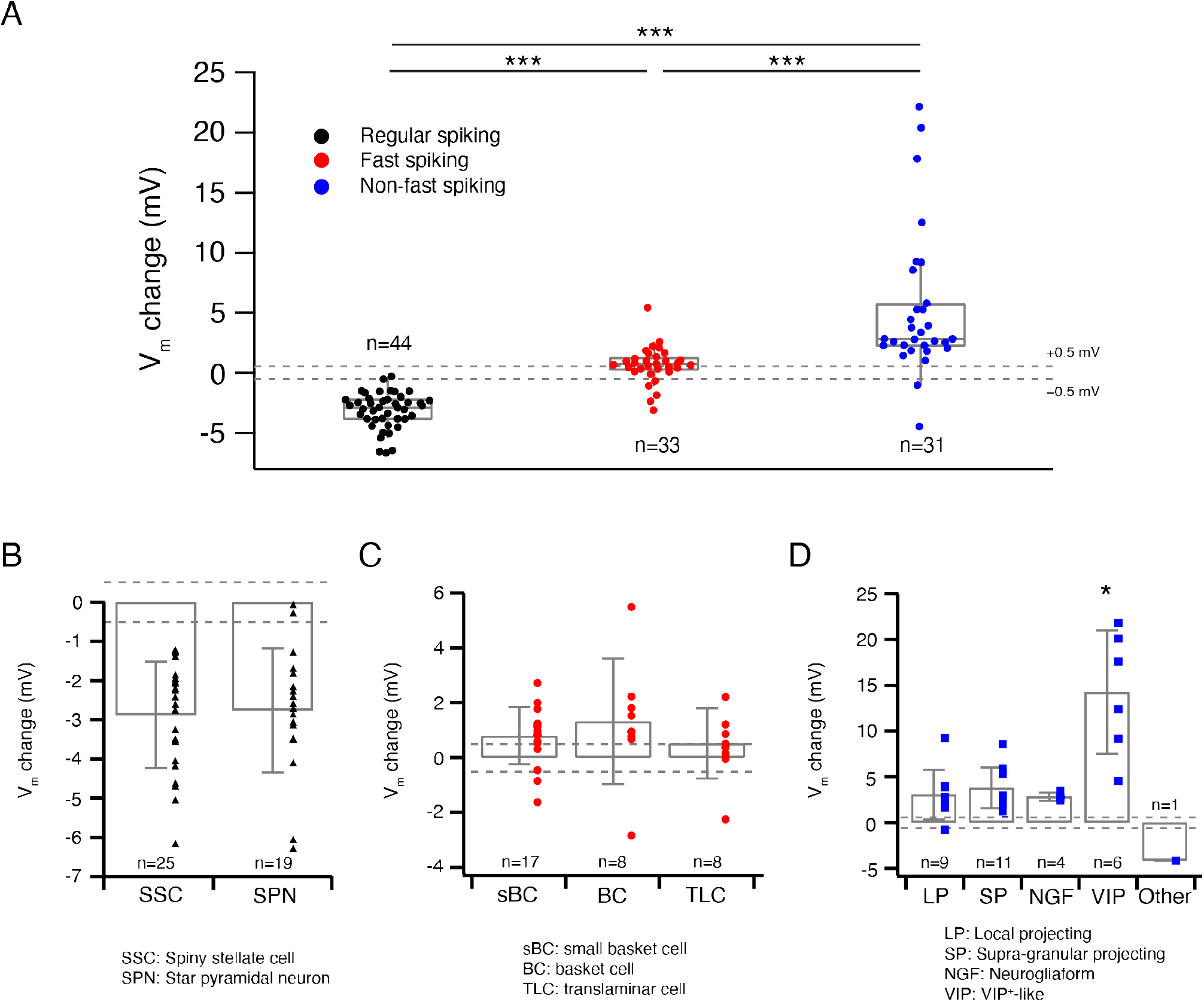
ACh-induced Vm changes are related to the L4 RS, FS and nFS neuron (sub)types. (A) Box plots of ACh-induced Vm changes in L4 RS, FS and nFS neurons. Individual data points are given on the right. P value was calculated using the non-parametric Wilcoxon-Mann-Whitney two-sample rank test. * p < 0.05, * * p < 0.01, * * * p < 0.001. Dashed lines indicate the Vm change at ± 0.5 mV. (B) Histograms of ACh-induced V_m_ changes for two L4 RS neuron subtypes: spiny stellate cells and star pyramidal neurons. No statistically significant difference was found between two subtypes. (C) Histograms of ACh-induced V_m_ changes for three L4 FS neuron subtypes: small basket cells, basket cells and translaminar cells. No statistically significant difference was found among three subtypes. (D) Histograms of ACh-induced V_m_ changes for five L4 nFS neuron subtypes: local projecting (putative SST+, non-Martinotti cell-like), supragranular projecting (putative SST+, Martinotti cell-like), NGF, VIP^+^-like and unclassified interneurons. VIP^+^-like interneurons show the strongest depolarisation of the five subtypes. Statistically significant differences were found between VIP+-like and three other interneuron subtypes (LP, SP, NG).

To evaluate the cell-type specificity of ACh-induced changes in V_m_ in more detail, we grouped the ACh response with respect to the L4 neuron subtype identified by the electrophysiological and morphological features described above. The two L4 RS excitatory neuron subtypes did not exhibit a significantly different ACh response (SSCs: -2.9 ± 1.4 mV, n = 25; SPNs: -2.8 ± 1.6 mV, n = 19; p = 0.85; **Fig. 3B**). Similarly, no significant difference was found in the ACh-induced change in V_m_ between three L4 FS interneuron subtypes (sBCs: 0.8 ± 1.0 mV, n = 17; BCs: 1.3 ± 2.3 mV, n = 8; TLCs: 0.5 ± 1.3 mV, n = 8; p = 0.42; **Fig. 3C**). In contrast, in four subtypes of L4 nFS interneurons the ACh-induced V_m_ change in VIP interneurons is significantly larger than that in the other three subtypes (LPs: 3.0 ± 2.7 mV, n = 9; SPs: 3.8 ± 2.2 mV, n = 11; NGFs: 2.8 ± 0.5 mV, n = 4; VIPs: 14.3 ± 6.7 mV, n = 6; p = 1.5 × 10_-3_) (**Fig. 3D**). We did not record the ACh response of the transcolumnar-projecting L4 nFS interneuron due to their scarcity.

### ACh differentially changes the intrinsic excitability of L4 FS and nFS interneurons

ACh not only modulates V_m_ but induces also changes in other intrinsic properties of L4 neurons. We have previously studied the effects of 100 µM ACh on the intrinsic properties of L4 excitatory neurons *(Eggermann and Feldmeyer 2009)* and demonstrated that ACh reduces their excitability through a hyperpolarisation of V_m_ and a reduction in R_in_. In this study, we focussed mainly on L4 interneurons. Low concentrations of ACh (30 µM) induced no significant change in the intrinsic properties of L4 FS interneurons except for the V_m_ (cf. **Figs. 2,3**). For example, no change was found for the AP half-width (Control: 0.26 ± 0.06 ms, n = 8; ACh: 0.26 ± 0.05 ms, n = 8; p = 0.31) and the AP amplitude (Control: 88.0 ± 12.6 mV, n = 8; ACh: 80.0 ± 8.9 mV, n = 8; p = 0.08) (**Fig. 4A and Table S1**). In contrast, apart from V_m_ changes (cf. **Figs. 2,3**) ACh also altered three other intrinsic electrophysiological properties of L4 nFS interneurons: the AP half-width was increased (Control: 0.44 ± 0.10 ms, n = 10; ACh: 0.48 ± 0.11 ms, n = 10; p = 0.04) and the AP amplitude was decreased (Control: 92.8 ± 10.9 mV, n = 10; ACh: 82.6 ± 11.4 mVs, n = 10; p = 2.0 × 10_-3_) (**Fig. 4B**). Furthermore, the rheobase current was significantly reduced by ACh (Control: 158.08 ± 75.5 pA, n = 10; ACh: 84.0 ± 113.1 pA, n = 10; p = 5.9 × 10_-3_) (**Table S1**). Thus, in contrast to L4 excitatory neurons, ACh enhanced the excitability of all recorded L4 nFS interneuron types.

**Figure 4.**
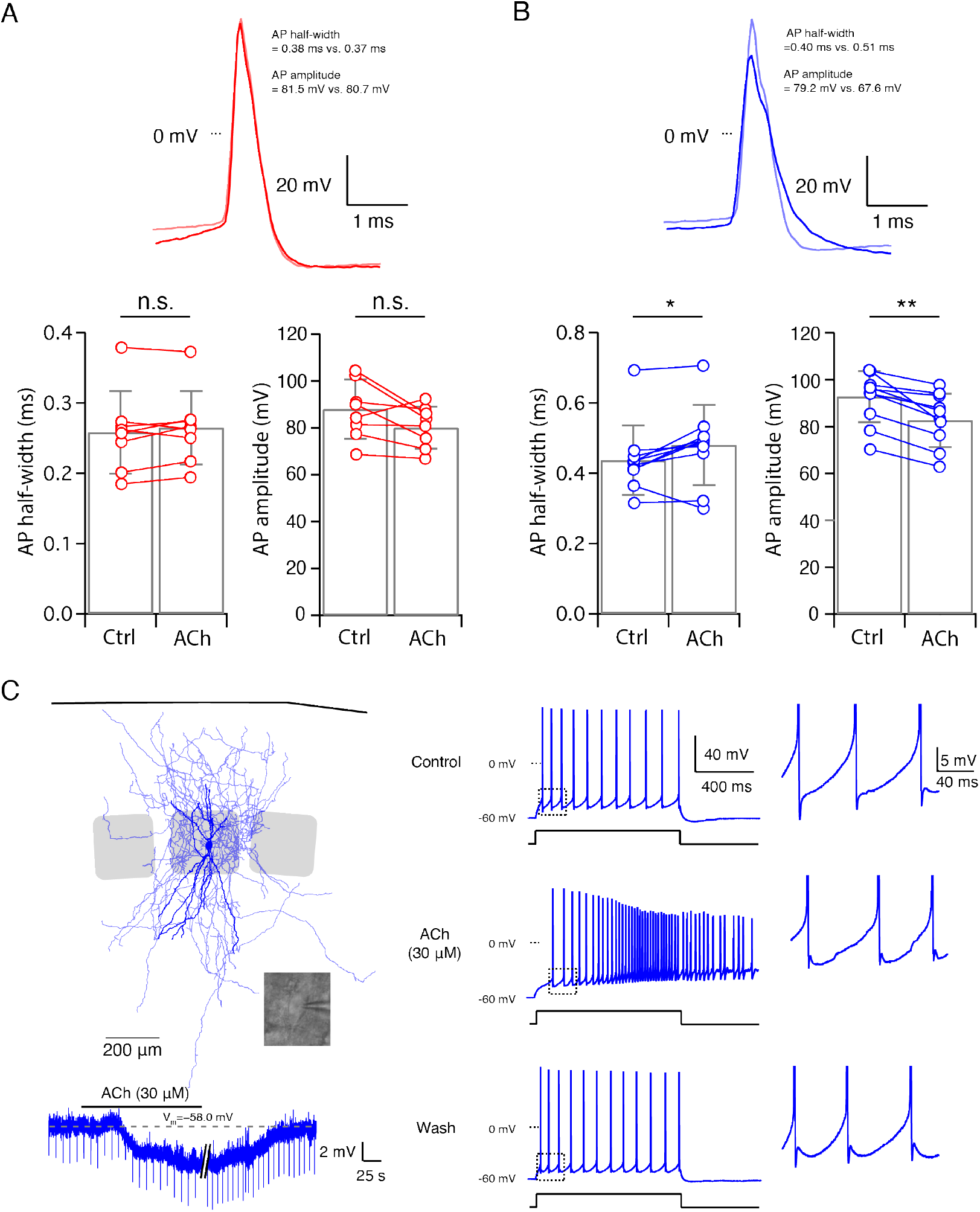
ACh-induced changes in other intrinsic properties of L4 FS and nFS interneurons. (A) Top, overlay of single APs recorded in a L4 FS interneuron before (light red) and after (red) the ACh application. Bottom, comparison histograms for AP half-width and AP amplitude. No statistically significant difference was found. (B) Top, overlay of single APs recorded in a L4 nFS interneuron before (light magenta) and after (magenta) ACh application. Bottom, comparison histograms for AP half-width and AP amplitude. A statistically significant increase in AP half-width and a decrease in AP amplitude were found. P value was calculated using the non-parametric Wilcoxon signed rank test. * p < 0.05, * * p < 0.01, * * * p < 0.001. (C) Example recording from a L4 nFS interneuron; soma and dendrites are in opaque, the axon in half-transparent blue. ACh induced an abnormal Vm in this neuron. Furthermore, a dramatic change in the firing pattern was found during the ACh application.

One particular L4 nFS interneuron (**Fig. 4C**) responded to the ACh application with a V_m_ hyperpolarisation, in contrast to most of L4 nFS interneurons. Furthermore, ACh changed its repetitive firing property (**Fig. 4C**). Under control condition, this neuron showed a regular spiking firing pattern with a small spike-frequency adaptation, which in the presence of 30 µM ACh was transformed to an accelerating firing pattern together with a spike amplitude accommodation. In addition, AP firing persisted even after terminating current injection. Because firing pattern and V_m_ returned to normal after washout (**Fig. 4C**), the marked alteration in the firing pattern cannot be the result of deteriorating recording conditions. Hence, already at low concentrations, ACh can dramatically change the electrophysiological behaviour of some L4 nFS interneurons.

### ACh-induced Vm changes in L4 neurons are mainly regulated by muscarinic receptors

To reveal the molecular mechanism of ACh-induced V_m_ changes in L4 neurons, slices were superfused with the general mAChR antagonist ATRO (200 nM) before application of ACh. A comparison of the ACh-induced change in V_m_ before and during co-application of ATRO showed that the mAChR antagonist completely blocks the response in all L4 RS excitatory neurons (Control: -3.8 ± 0.9 mV, n = 9; ATRO: -0.4 ± 0.5 mV, n = 9; p = 3.9 × 10_-3_) (**Fig. 5A,D**) and all L4 FS interneurons (Control: 1.5 ± 0.6 mV, n = 4; ATRO: 0.1 ± 0.3 mV, n = 4; p = 0.13) (**Fig. 5B,D**). This suggests that the ACh-induced V_m_ changes in these L4 neuron types are exclusively mediated by mAChRs. In contrast, in L4 nFS interneurons, ATRO largely (but not completely) blocks the V_m_ change induced by 30 µM ACh (Control: 9.2 ± 7.2 mV, n = 11; ATRO: 3.7 ± 4.5 mV, n = 11; p = 9.8 × 10^−4^) (**Fig. 5C,D**). In the majority (8 out of 11) of L4 nFS interneurons, atropine nearly completely blocked the ACh-induced V_m_ change while in the remainder (3 out of 11), a residual ACh-induced change in V_m_ still exists after the co-application of ATRO. We tested whether this residual depolarisation was mediated by nAChRs (see below). To identify the mAChR type mediating the modulatory effect, TRO (1 µM), a specific M4 mAChR antagonist, and PIR (0.5 µM), a specific M1 mAChR antagonist, were applied before ACh. We found that TRO completely blocked the ACh-induced hyperpolarisation in L4 RS excitatory neurons (**Fig. S3A**) while PIR completely blocked the ACh-induced depolarisation in L4 FS interneurons (**Fig. S3B**). However, in L4 nFS interneurons, PIR blocked the ACh-induced depolarisation only partially (**Fig. S3C**).

**Figure 5.**
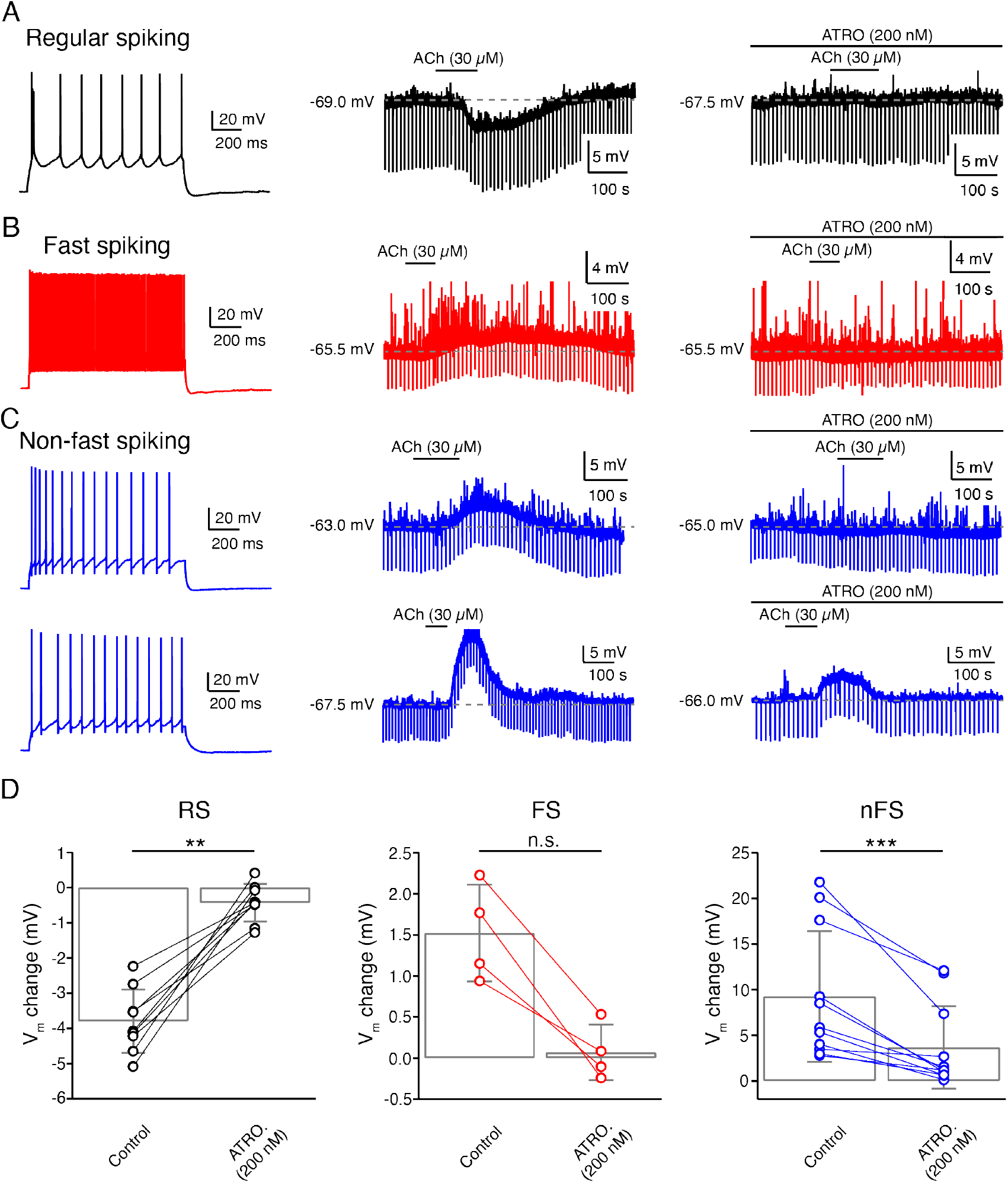
ACh-induced V_m_ changes in L4 RS, FS and nFS neurons are mainly regulated by muscarinic ACh receptors. (A) An example recording of the time course of ACh-induced Vm change under control condition and in the presence of atropine, a general mAChR antagonist, in a L4 RS neuron. (B) Same as (A) but for a L4 FS interneuron. (C) Same as (A,B) but for two L4 nFS interneurons: upper, sub-threshold depolarisation which could be completely blocked by atropine; lower, supra-threshold depolarisation which could be partly blocked by atropine. (D) Comparison histograms for L4 RS (left), FS (middle) and nFS (right) neurons under control conditions and in the presence of atropine. P value was calculated using the non-parametric Wilcoxon signed rank test. * p < 0.05, ** p < 0.01, *** p < 0.001, n.s. p >= 0.05.

### L4 VIP+-like nFS interneurons are strongly depolarised by low-concentration of ACh via both muscarinic and nicotinic receptors

The fact that a subset of L4 nFS interneurons showed a strong ACh-induced depolarisation (**Fig. 2C and Fig. 3A,D**) that was not fully blocked by ATRO (**Fig. 5C,D**) indicates that these interneurons may respond to ACh via both mAChRs and nAChRs. In those L4 nFS interneurons, in which ATRO blocked the ACh-induced depolarisation only incompletely, MEC (1 µM), a general nAChR antagonist, together with ATRO were applied before ACh. An example recording from a putative VIP+ L4 nFS interneuron (**Fig. 6A**) is shown in Fig. 6B. This neuron exhibits an irregular firing pattern, a bipolar dendritic structure, and a narrow translaminar axonal projection, all of which are characteristics typical of VIP+ interneurons *(Porter et al. 1998; Pronneke et al. 2015; Emmenegger et al. 2018)*. ACh induced a strong depolarisation in this neuron and elicited spontaneous AP firing. Even in the presence of ATRO, the ACh-induced AP firing still exists. Only when ATRO and MEC were applied together, the ACh-induced change in V_m_ was blocked (**Fig. 6B**). A nAChR-mediated depolarisation was observed not only in L4 VIP+ interneurons (n = 2) but also in one putative somatostatin-positive (SST+) interneuron and one neurogliaform (NGF) cell (**Fig. 6C**). However, only VIP+ interneurons showed such a strong nAChR-mediated depolarisation. In one recording from a L4 VIP+ interneuron, we found that the ATRO-resistant depolarisation was completely blocked by DHβE, a specific antagonist for α4β2-subunit containing nAChRs (**Fig. S3C**).

**Figure 6.**
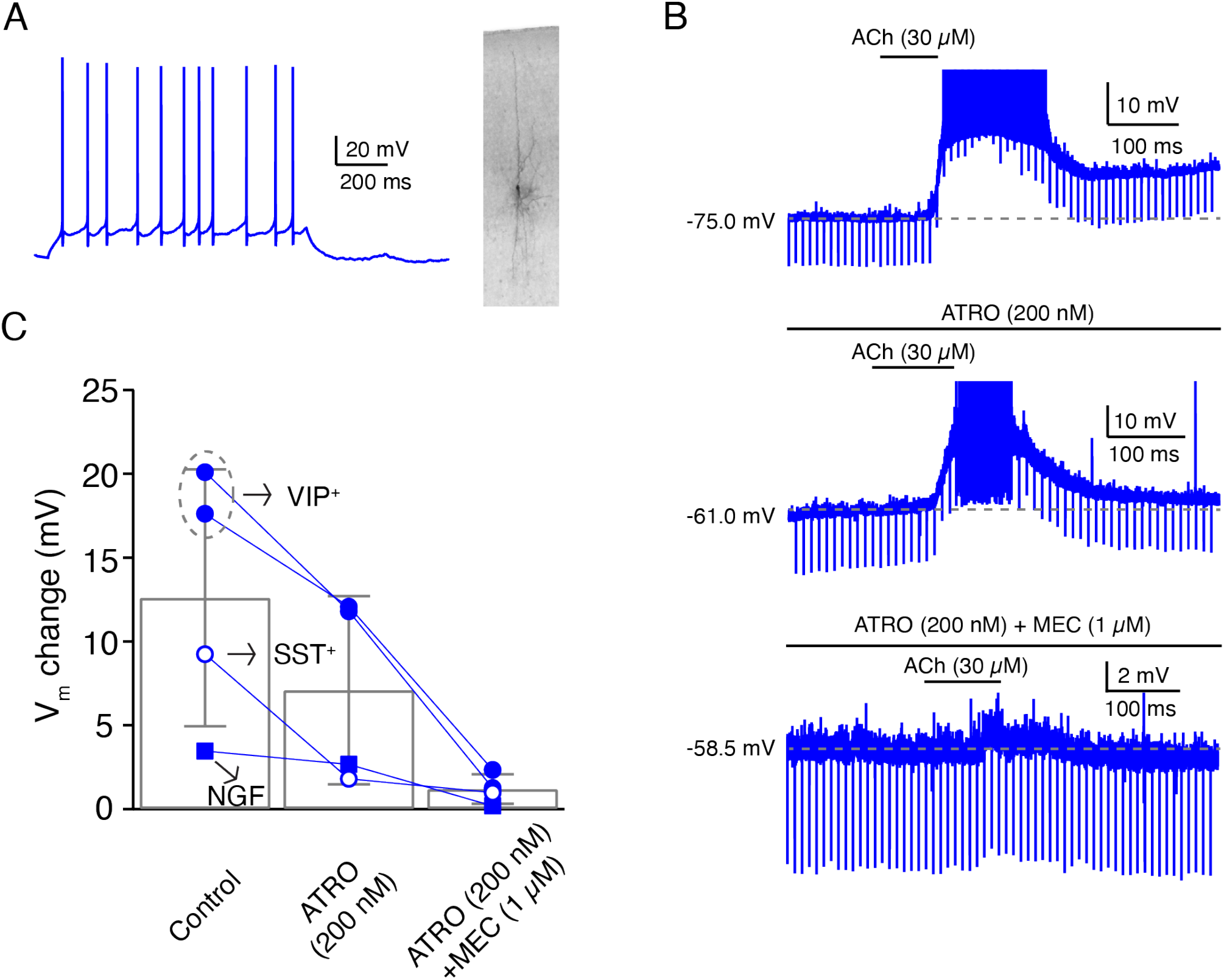
A subpopulation of L4 non-FS interneurons are strongly depolarised by ACh via both muscarinic and nicotinic ACh receptors. (A) Firing pattern (left) and morphology (right) of an example L4 VIP^+^-like interneuron. (B) Time course of V_m_ change during ACh application alone (top), during the application of ACh in the presence of atropine (middle) and during the ACh application in the presence of atropine and mecamylamine (bottom). (C) Histogram of ACh-induced V_m_ changes under three conditions. Note that in these four L4 nFS interneurons, two are VIP+-like, one is SST+-like and one is a NGF cell.

## Discussion

In the present study, we found that all L4 neurons are persistently modulated by low concentrations of ACh in a cell-type specific way (see **Fig. 7**): (1) ACh (30 µM) reduces the intrinsic excitability of L4 RS excitatory neurons by activating the M4 mAChRs presumably located in the soma and/or dendrite, which leads to a hyperpolarisation of V_m_ and a decreased R_in_; (2) ACh induces a small but significant depolarisation in L4 FS interneurons by activating the M1 mAChRs; (3) ACh elicits a markedly stronger depolarisation in L4 nFS interneurons compared to L4 FS interneurons by activating not only mAChRs (of the M1 and/or M3/5 type) but also nAChRs (presumably of the α4β2* type); (4) In a subset of L4 nFS interneurons, the VIP+-like interneurons, the ACh-induced depolarisation was sufficiently large to induce spontaneous AP firing through activation of both mAChRs and nAChRs.

**Figure 7.**
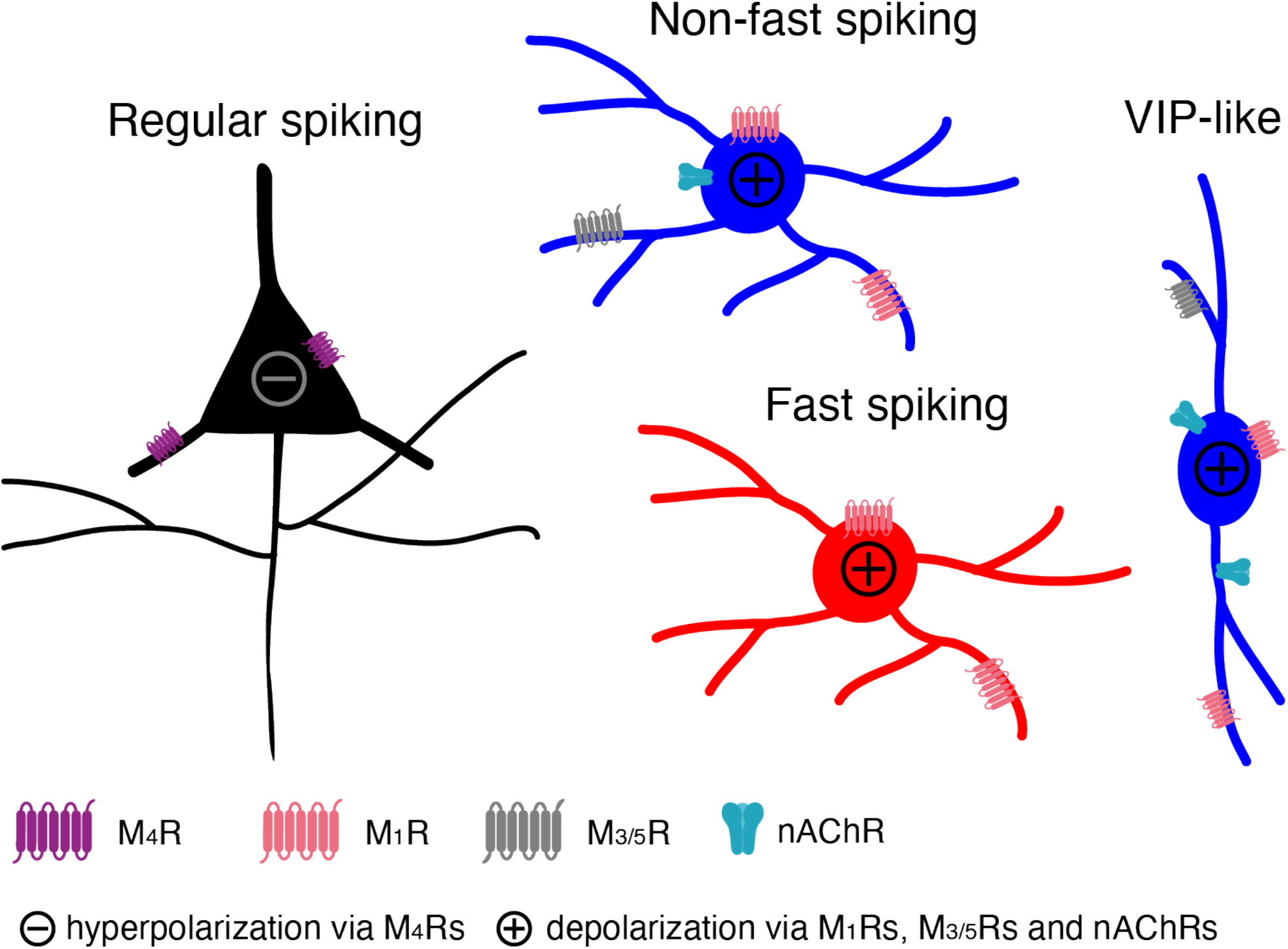
A cartoon summarising the modulatory effects of low concentrations ACh on L4 RS, FS and nFS neurons and their potential molecular mechanisms.

### L4 neuronal cell-type classification

A detailed neuronal cell-type classification is necessary and critical an in-depth understanding the modulatory effects of ACh. Traditionally, neurons are classified based on their morphological (dendritic and axonal structures) and electrophysiological (repetitive firing behaviours) properties *(Petilla Interneuron Nomenclature et al. 2008; DeFelipe et al. 2013)*. With the development and sophistication of single-cell mRNA sequencing techniques, the molecular features of neurons add an additional layer of complexity to neuronal classification *(Zeng and Sanes 2017; Yuste et al. 2020)*. We have performed a series of studies to dissect the neuronal diversity in layer 4 of rat barrel cortex *(Feldmeyer et al. 1999; Lubke et al. 2000; Koelbl et al. 2015; Emmenegger et al. 2018)*. In general, layer 4 comprises three neuronal cell classes showing distinct repetitive firing properties: regular spiking, fast spiking and non-fast spiking (adapting, irregular, late etc). Taking the morphological diversity also into account, L4 RS excitatory neurons have been classified into spiny stellate cells and star pyramidal neurons *(Feldmeyer et al. 1999; Lubke et al. 2000)* while L4 FS interneurons have been divided into cluster 3 (small basket cells), cluster 2 (basket cells) and translaminar-projecting FS interneurons *(Koelbl et al. 2015)*. Most of these L4 FS interneurons are parvalbumin-positive but calbindin-negative. L4 nFS interneurons, on the other hand, have been separated into five morpho-electrophysiological subtypes including transcolumnar-projecting interneurons with an adapting firing pattern, locally projecting with an adapting firing pattern (presumably non-Martinotti cells), supragranular-projecting with an adapting firing pattern with a Martinotti-cell appearance, VIP+ cell-like with an irregular firing pattern (VIP+-like) and neurogliaform cells *(Emmenegger et al. 2018)*. The former three subtypes are somatostatin-positive while the latter two Prox1-positive. Our classification of L4 neurons is in line with several other groups focusing on the barrel cortex or primary visual cortex of rats and mice *(Gibson et al. 1999; Porter et al. 2001; Beierlein et al. 2003; Ma et al. 2006; Scala et al. 2019)*.

### The necessity of bath-application of low-concentration ACh to study the tonic neuromodulation mediated by muscarinic receptors

ACh modulates the intrinsic neuronal properties through two categories of cholinergic receptors, i.e. mAChRs and nAChRs. mAChRs are G-protein coupled receptors the activation of which initiates a signalling cascade inside the neuron. In contrast, nAChRs form ligand-gated cation channels *(Unwin 2003; Dani 2015)*. These two types of receptors work at different concentrations of ACh. mAChRs already show a high affinity to ACh at low concentrations (in the range of 1-100 µM) while nAChRs require a high concentration of ACh for channel opening (in the mM range). Previous studies have used puff-application of 1-10 mM ACh to study the nicotinic effects of ACh on excitatory and inhibitory neurons in several cortical areas *(Xiang et al. 1998; Gulledge and Stuart 2005; Gulledge et al. 2007; Poorthuis et al. 2013b; Poorthuis et al. 2013a)*. Puff-application of agonists has a high spatiotemporal resolution and is a suitable strategy to simulate the phasic ACh release at cholinergic presynaptic terminals. However, at such high concentrations, ACh will not be hydrolysed (and hence inactivated) immediately so that it may persist at low concentrations in the perisynaptic space. Bath application of an agonist is a good approach to simulate the latter condition because the agonist concentration will be maintained at a constant level to allow the measurement of neuronal properties at equilibrium. In the extracellular space, it is likely that a neurotransmitter/neuromodulator is only present at µM concentration because of its rapid diffusion from its release site *(Borroto-Escuela et al. 2015)*. In addition, application of high concentrations of ACh will mask the effects mediated by mAChRs so that the application of low concentrations (∼µM) of ACh is a prerequisite to uncover their functional effects.

### Unique ACh modulation of L4 excitatory neurons

We have shown previously that 100 µM ACh persistently hyperpolarises L4 excitatory neurons and reduces their intrinsic excitability *(Eggermann and Feldmeyer 2009)*, an ACh effect markedly different from that on most of pyramidal cells except for L6A corticocortical neurons *(Yang et al. 2020)*. In pyramidal cells, ACh induces a depolarisation and therefore enhances the excitability. The ACh effects on L4 excitatory neurons are exclusively mediated by mAChRs. Similar findings were obtained in another study using optogenetic activation of synaptic ACh release *(Dasgupta et al. 2018)*. Here, using a lower concentration of ACh (30 µM), similar results were obtained. Note that, as discussed above, 30 µM is an ACh concentration closer to the physiological range than 100 µM. Through bioctyin labelling, the morphology of the recorded L4 RS excitatory neurons was also revealed. When comparing the ACh-induced changes in V_m_ for the two L4 RS subtypes, no clear difference was found indicating that the ACh response in both L4 RS subtypes is similar.

### ACh persistently depolarises L4 FS interneurons

The effects of ACh on FS interneurons has been a long-standing matter of debate. Conflicting results have been published by different research groups. Puff-application of 5 mM ACh induced a transient hyperpolarisation that was mediated by mAChRs in rat neocortical L5 FS interneurons *(Xiang et al. 1998)*. Recently, in the mouse visual cortex it has been shown that optogenetically stimulated ACh release led to an indirect inhibition in L2/3 FS interneurons via ‘facilitation’ of the cholinergic responses in L2/3 somatostatin-positive interneurons (*Chen et al. 2015*). On the other hand it has been postulated that ACh does not affect the V_m_ of FS interneurons. In the rat frontal cortex, bath-application of carbachol (10 µM), a cholinergic agonist, had no effect on L2/3 FS interneurons *(Kawaguchi 1997)*. In a follow-up study the same group used focal application of ACh (100 µM or 5 mM for comparison with the study by *Xiang et al*., *1998*) onto FS interneurons in rat visual and prefrontal cortex; the authors concluded that the focal application itself (i.e. a mechanical artefact but not the transient ACh exposure) caused the hyperpolarising response *(Gulledge et al. 2007)*. Except for *Xiang et al. (2018)*, most investigators have shown that there is no direct effect of ACh on FS interneurons *(Muñoz and Rudy 2014)*; however they reported a presynaptic effect of ACh. In contrast to previous studies, we found a persistent ACh-induced V_m_ depolarisation in L4 FS interneurons, an effect that appeared in all three subtypes of L4 FS interneurons. To the best of our knowledge, this is the first time that a direct depolarising effect of ACh on FS interneurons has been demonstrated conclusively. In addition, we were able to show that this effect is mediated by M1 mAChRs. The ACh-induced depolarisation persisted in the presence of GABA and glutamate receptor antagonists so that indirect effects of ACh can be excluded. We were unable to investigate the effect of ACh on another subtype of FS interneurons, the chandelier or axo-axonic cells, which are very scarce if not absent in cortical layer 4 *(Wang et al. 2019)*.

### ACh modulates L4 nFS interneuron activity in a subtype-specific way

L4 nFS interneurons are a heterogenous population that show diverse firing pattern, dendritic/ axonal morphology and molecular expression pattern. To elucidate modulatory effects of ACh on L4 nFS interneurons, a clear separation into identifiable subtypes is required. The local- and supragranular-projecting subtypes of L4 nFS interneurons display an adapting firing pattern similar to that of somatostatin-positive (SST+) interneurons. Indeed, immunocytochemistry revealed that both subtypes of L4 nFS interneurons are somatostatin-positive *(Emmenegger et al. 2018)*. Here, we found that L4 SST+-like interneurons including both local projecting (non-Martinotti-like) and supragranular projecting (Martinotti-like) cells responded to ACh with a strong depolarisation that is predominantly mediated by mAChRs. Consistent with our findings, in the mouse barrel cortex it has been shown that L4 SST+ interneurons were depolarised and fired spikes in response to bath-applied muscarine (3 µM) *(Xu et al. 2013)*. In one L4 SST+-like interneuron, ACh induced a depolarisation mediated by both mAChRs and nAChRs suggesting that even at µM ACh concentrations activation of nAChRs may be also possible. In a previous study, it has been shown that ACh directly excites SST+ neurons via both mAChRs and nAChRs in layer 2/3 of mouse visual cortex *(Chen et al. 2015)*. However, a very high concentration of ACh (10 mM) was puff-applied in that study, which is very different from the bath-application of ACh (30 µM) described here.

Excitation of VIP+ interneurons by nAChRs has been observed in several cortical areas of both rat and mouse *(Porter et al. 1999; Férézou et al. 2007; Koukouli et al. 2017; Askew et al. 2019; Prönneke et al. 2020)*. In the rat motor cortex, local pressure application of ACh (100 µM) or the selective nAChR agonist DMPP (100–500 mM) depolarised VIP+ interneurons located in layer 3-5 and induced a discharge of action potentials *(Porter et al. 1999)*. Pharmacological experiments suggested that the ACh effect was mediated by non-α7 nicotinic receptors containing α4β2 and α5 subunits *(Koukouli et al. 2017)*. In another study from the same group, it was shown that bath-application of nicotine (1 µM) also resulted in a strong depolarisation leading to a sustained action potential discharge in VIP+ interneurons *(Férézou et al. 2007)*. Similarly, bath-application of nicotine (1 µM) in the mouse auditory cortex caused sustained AP discharge in VIP+ interneurons across the layers *(Askew et al. 2019)*. In the mouse barrel cortex, bath-application of ACh (40 µM) efficiently depolarised L2/3 VIP+ interneurons and changed the firing pattern from bursting to tonic spiking in a subpopulation *(Prönneke et al. 2020)*; however, the authors concluded that cholinergic modulation was mediated exclusively by nAChRs. All of the aforementioned studies emphasised the critical role of nAChRs in the cholinergic modulation of VIP+ interneurons but overlooked any direct involvement of mAChRs. However, our recordings from L4 VIP+ interneurons demonstrated that both AChR types participate in the cholinergic modulation in a cooperative way because ATRO partially blocked the depolarisation or shortened the duration of repetitive action potential firing induced by ACh; ATRO together with nAChR antagonist MEC completely blocked the ACh effect. Similarly, in rat frontal cortex, VIP+ cells showed a sustained V_m_ depolarisation in response to bath-applied muscarine (3 µM) in the presence of TTX *(Kawaguchi 1997)* indicating that mAChRs are expressed in these interneurons.

For NGF cells, the focus of attention is mostly cortical layer 1 where NGF cells are abundant *(Christophe et al. 2002; Gulledge et al. 2007; Arroyo et al. 2012; Brombas et al. 2014)*. Puff-application of nicotinic agonists such as ACh, DMPP, choline onto L1 NGF cells or optogenetic stimulation of cholinergic fibres in layer 1 has revealed nicotinic excitation of NGF cells. Similar results have been found in L2/3 5-HT_3a_R+ NGF cells of mouse barrel cortex *(Lee et al. 2010)*. Here, we found that in L4 NGF cells of the barrel cortex, low concentrations of ACh led to a mAChR-mediated sustained depolarisation. In one L4 NGF cell, a participation of nAChRs in depolarisation was also found.

In addition to SST+, VIP+ and NGF cells, layer 4 comprises other nFS subtypes *(Tasic et al. 2018)*. In one L4 nFS interneuron, we were able to show that, in contrast to most other L4 nFS interneurons, ACh application resulted in a hyperpolarisation of this neuron and dramatically changed its repetitive firing pattern during the suprathreshold current injection. From the cholinergic response of this neuron together with its firing pattern and morphology, it is reminiscent of a subset of cholecystokinin-positive (CCK+) cells in L2/3 of rat frontal cortex which exhibited a prominent hyperpolarisation in response to muscarine (3 µM) and had large somata and extensive axonal arbors *(Kawaguchi 1997)*. Therefore, the nFS interneuron showing the ACh-induced hyperpolarisation could be a L4 CCK+ cell. Similarly, some hippocampal CA1 CCK+ interneurons showed also an ACh-induced hyperpolarisation mediated by mAChRs *(McQuiston and Madison 1999a; Cea-del Rio et al. 2011)*. Furthermore, the dramatic change in firing pattern induced by ACh has also been demonstrated in hippocampal CA1 CCK+ interneurons *(McQuiston and Madison 1999b; Lawrence et al. 2006; Cea-del Rio et al. 2010; Cea-del Rio et al. 2011)*. In these neurons, through the activation of M1 and M3 mAChRs the AHP was replaced by an afterdepolarisation which is often large enough to evoke APs in the absence of further stimulation *(McQuiston and Madison 1999b; Cea-del Rio et al. 2011)*.

### Functional significance of cholinergic neuromodulation for L4 neuronal microcircuits

In the neocortex, the ACh is continuously released to the extracellular space and its levels change dramatically during different behavioural states *(Teles-Grilo Ruivo et al. 2017)*. Most of the intracortical ACh is not released at synaptic contacts but rather diffusely into the extracellular space through an extrasynaptic volume transmission *(Fuxe and Borroto-Escuela 2016)*. Under this condition, cholinergic modulation is spatiotemporally slower but broader, thereby tuning neuronal network function. Modulation of neurons and their synaptic interactions through mAChRs may induce neuronal oscillations and therefore change the information processing mode in L4 neuronal microcircuits. Indeed, it has been demonstrated that bath-application of carbachol (10 µM) to activate mAChRs and kainate (300 nM) to increase the tonic excitatory drive elicited persistent gamma frequency network oscillations in cortical layer 4 of mouse barrel cortex *(Buhl et al. 1998)*. In addition, differential modulation of L4 excitatory and inhibitory neurons, i.e. the persistent hyperpolarisation of L4 excitatory neurons and depolarisation of L4 inhibitory neurons, will change the excitation-inhibition balance towards inhibition and reduce the responsiveness of the L4 recurrent excitatory microcircuit. Therefore, our results support the hypothesis that ACh has a filtering action in the major recipient layer of the neocortex *(Eggermann and Feldmeyer 2009)*. Because neocortical layer 4 is uniquely positioned to gate thalamocortical input to the neocortex, cholinergic modulation of L4 neuronal microcircuits will affect the whole barrel cortex together with the related cortical areas (e.g. M1 and S2) and finally the animal behaviour *(Eggermann et al. 2014; Meir et al. 2018)*. In addition, our finding that mAChRs ubiquitously but differently modulate the activity of L4 excitatory and inhibitory neurons might open the door to more specific therapeutic strategies to treat cognitive dysfunction or psychiatric disorders linked to degeneration of the cholinergic system in diseases such as such as Alzheimer’s disease and schizophrenia *(Marin 2012; Hampel et al. 2018)*.

## Author contributions

GQ designed research, performed experiments, analyzed data, and wrote the draft manuscript. DF supervised the work and wrote the manuscript. All authors contributed to the article and approved the submitted version.

## Funding

This work was supported by the Helmholtz Society, the DFG Research Group - BaCoFun (grant no. Fe471/4-2 to D.F.); European Union’s Horizon 2020 Research Innovation Programme (grant agreement no. 785907; HBP SGA2 to D.F.).

## Acknowledgements

We are grateful to Dr Danqing Yang for insightful comments and suggestions on the manuscript. We thank Werner Hucko for excellent technical assistance and Dr Karlijn van Aerde for custom-written macros in Igor Pro software. We also thank Dr. Vishalini Emmenegger for help with Neurolucida reconstructions.

## Notes

### Competing Interest Statement

The authors have declared no competing interest.

