## Supplementary Materials for "Cell-type specific neuromodulation of excitatory and inhibitory neurons via muscarinic acetylcholine receptors in layer 4 of rat barrel cortex"

### Summary

Supplementary materials include 3 figures and 1 table.

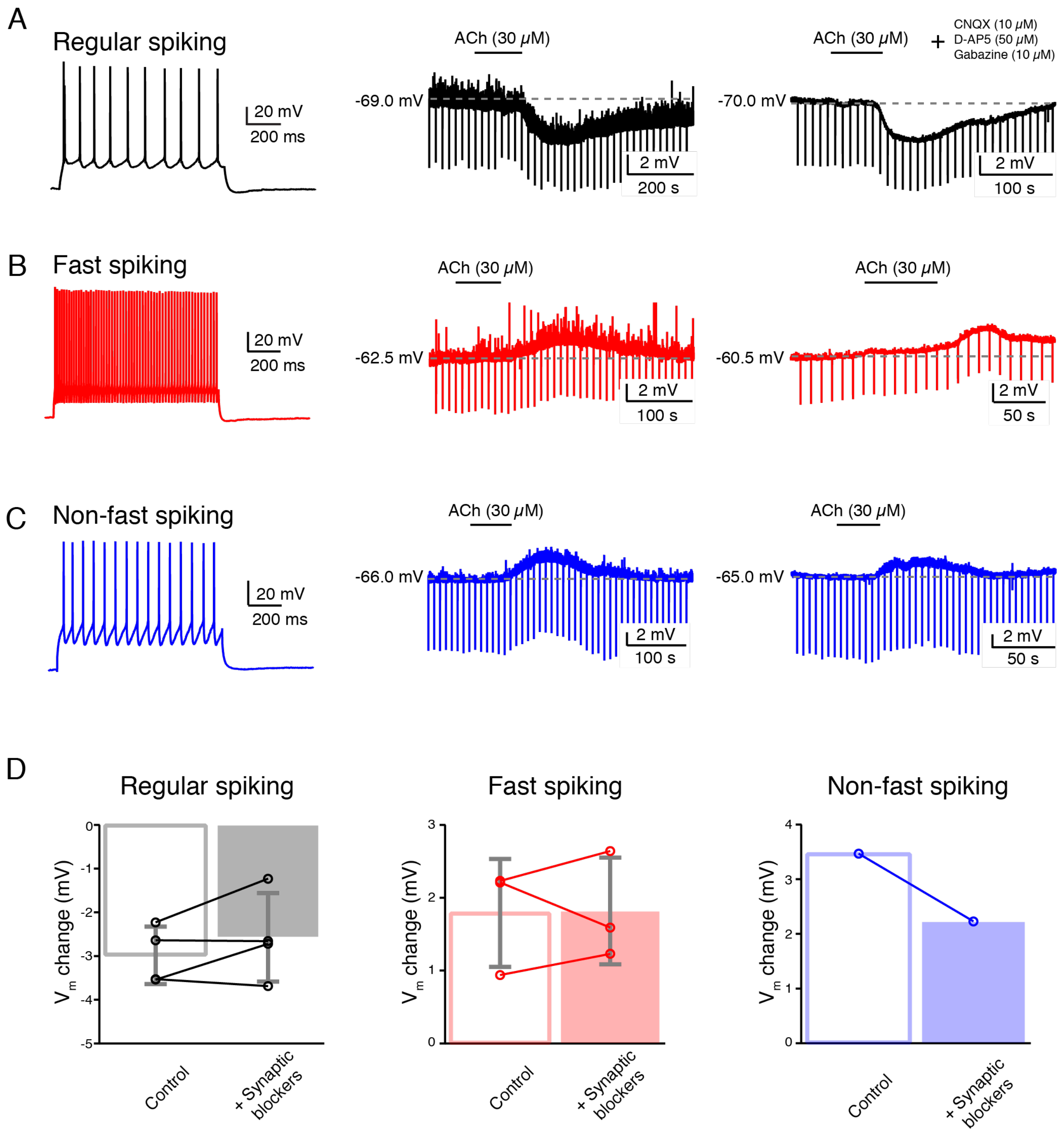


**Figure S1. ACh-induced Vm changes are independent of synaptic transmission.**

(A) An example recording of the time course of ACh-induced Vm change under control condition (middle) and in the presence of synaptic blocker cocktail in a L4 RS neuron.

(B) Same as (A) but for a L4 FS interneuron.

(C) Same as (A,B) but for a L4 nFS interneuron.

(D) Comparison histograms of ACh-induced Vm changes for L4 RS (left), FS (middle) and nFS (right) neurons. No statistically significant difference was found.

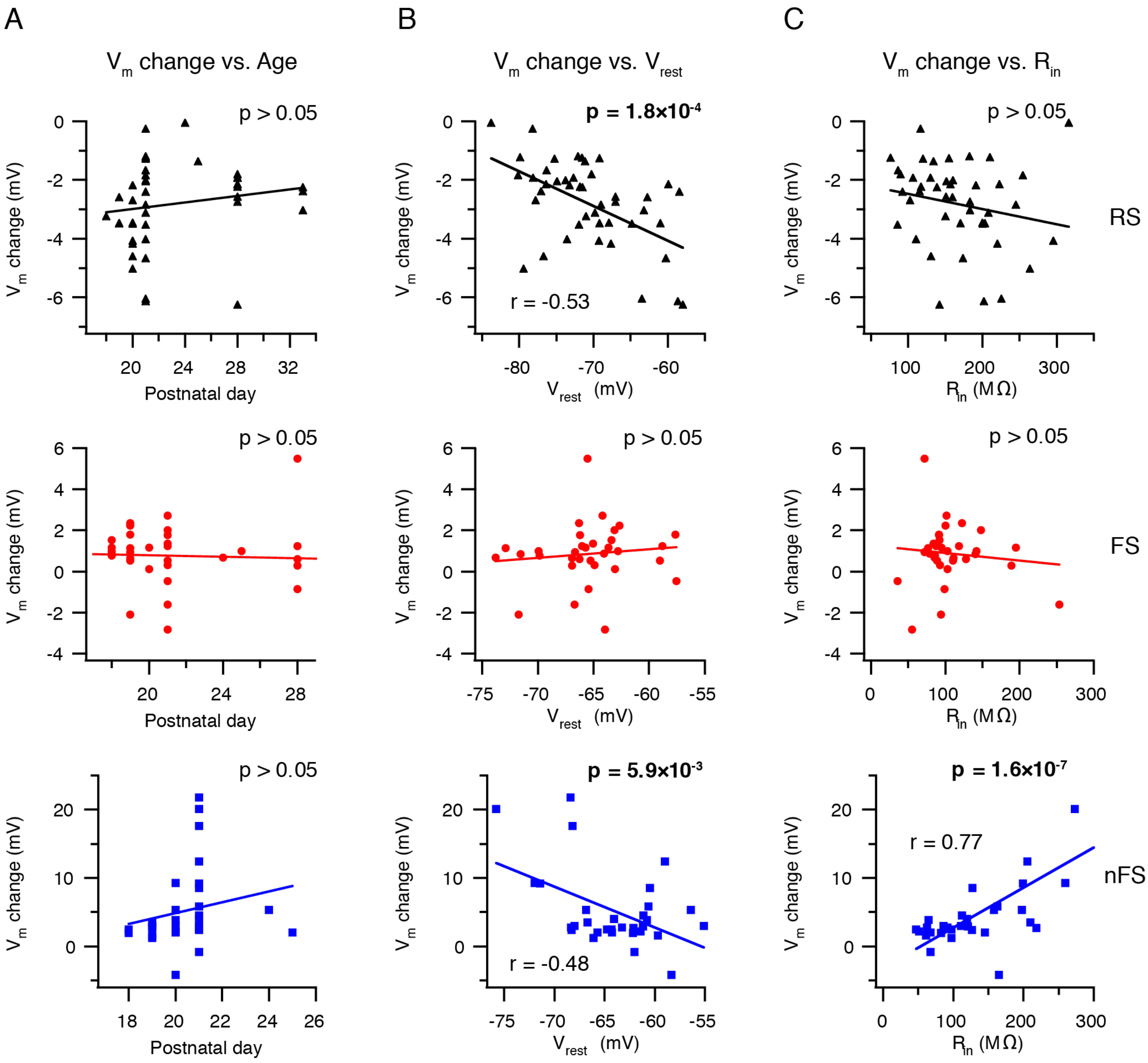


**Figure S2. Correlation analysis between the Vm change and the age, the resting membrane potential (Vrest) and the input resistance (Rin).**

(A) Correlation analysis between the Vm change and the age for L4 RS (top), FS (middle) and nFS (bottom) neurons. No statistically significant correlation was found.

(B) Same as (A) but between the Vm change and Vrest. Statistically significant correlations were found in L4 RS and nFS neurons.

(C) Same as (A,B) but between the Vm change and Rin. A statistically significant correlation was found in L4 nFS neurons.

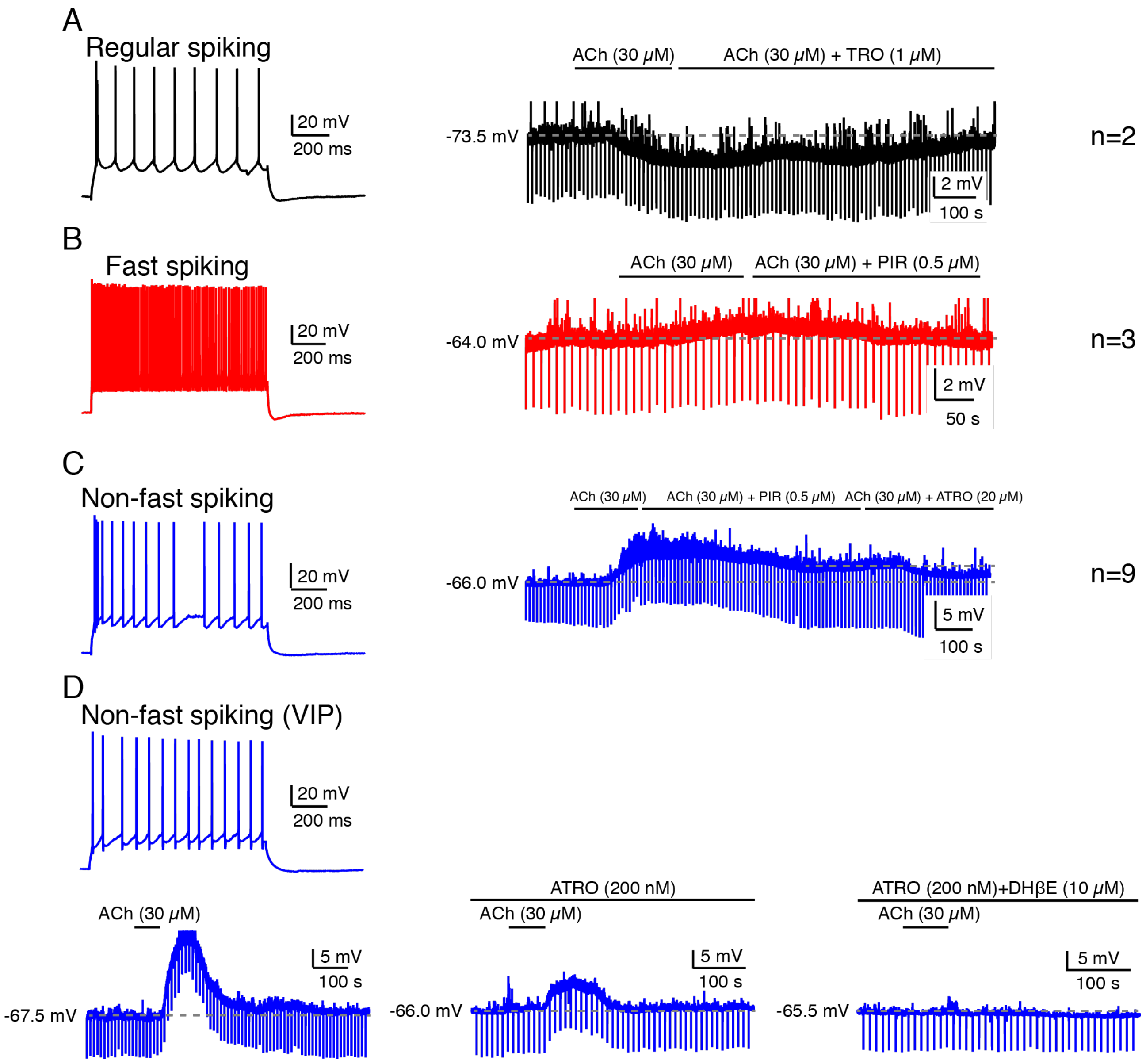


**Figure S3. Example recordings from L4 RS, FS and nFS neurons to unveil the subtype of muscarinic and nicotinic receptors that participate in the neuromodulation.**

(A) An example recording of the time course of Vm change during the ACh application and during the co-application of ACh and tropicamide, a specific M4 muscarinic receptor antagonist, in a L4 RS neuron. The number of recorded neurons is given on the right.

(B) An example recording of the time course of Vm change during the ACh application and during the co-application of ACh and pirenzepine, a specific M1 muscarinic receptor antagonist, in a L4 FS neuron.

(C) An example recording of the time course of Vm change during the ACh application, during the co-application of ACh and pirenzepine and during the co-application of ACh and atropine in a L4 nFS neuron. Note the residual Vm depolarization during the co-application of ACh and pirenzepine.

(D) An example recording of the time course of Vm change during the ACh application (lower left), during the co-application of ACh and atropine (lower middle) and during the co-application of ACh, atropine and dihydro-ß-erythroidine (lower right), a specific α4β2 subunit-containing nicotinic receptor antagonist, in a L4 nFS neuron.

**Table S1. ACh-induced changes in other intrinsic properties of L4 FS and nFS interneurons.**

P value was calculated using the non-parametric Wilcoxon signed rank test.

| | **L4 FS Ctrl** (n=8) | **L4 FS ACh** (n=8) | **L4 nFS Ctrl** (n=10) | **L4 nFS ACh** (n=10) | **p value** (**L4 FS Ctrl vs. ACh**) | **p value** (**L4 nFS Ctrl vs. ACh**) |
|---|---|---|---|---|---|---|
| ***Passive*** | | | | | | |
| Rin (MΩ) | 124.7 ± 64.3 | 123.8 ± 70.5 | 123.2 ± 71.2 | 136.8 ± 43.5 | 1,00 | 0,11 |
| Tau (ms) | 8.2 ± 1.5 | 9.3 ± 1.6 | 11.7 ± 4.9 | 12.3 ± 4.9 | 0,44 | 0,084 |
| Sag (%) | 7.6 ± 2.9 | 3.6 ± 6.5 | 11.5 ± 6.1 | 8.0 ± 7.9 | 0,44 | 0,28 |
| ***Single AP*** | | | | | | |
| Rheobase current (pA) | 258.3 ± 141.5 | 225.0 ± 113.8 | 158.0 ± 75.5 | 84.0 ± 113.1 | 0,41 | **$5.9\times10^{-3}$** |
| AP threshold (mV) | -31.6 ± 7.0 | -32.6 ± 7.0 | -36.4 ± 7.1 | -35.8 ± 5.7 | 0,74 | 0,49 |
| AP half-width (ms) | 0.26 ± 0.06 | 0.26 ± 0.05 | 0.44 ± 0.10 | 0.48 ± 0.11 | 0,31 | **0,037** |
| AP amplitude (mV) | 88.0 ± 12.6 | 80.0 ± 8.9 | 92.8 ± 10.9 | 82.6 ± 11.4 | 0,078 | **$2.0\times10^{-3}$** |
| AHP amplitude (mV) | 20.4 ± 2.6 | 20.6 ± 3.9 | 13.1 ± 4.5 | 12.7 ± 4.1 | 0,74 | 0,77 |
| ***Repetitive firing*** | | | | | | |
| Max. firing frequency (Hz) | 108.5 ± 49.5 | 126.9 ± 33.2 | 36.9 ± 11.8 | 39.8 ± 16.4 | 0,14 | 0,98 |
| Slope of F-I curve (APs/ 100 pA) | 37.2 ± 10.3 | 37.2 ± 8.6 | 16.1 ± 7.4 | 21.5 ± 18.1 | 0,74 | 1,00 |
